# HIV viral load and the efficacy of antiviral drug

**DOI:** 10.1101/2020.11.06.372094

**Authors:** Stephanie Yunfei Zhang, Mesfin Asfaw Taye

## Abstract

Developing antiviral drugs is an exigent task since viruses mutate to overcome the effect of antiviral drugs. As a result, the efficacy of most antiviral drugs is short-lived. To include this effect, we modify the Neumann and Dahari model. Considering the fact that the efficacy of the antiviral drug varies in time, the differential equations introduced in the previous model systems are rewritten to study the correlation between the viral load and antiviral drug. The effect of antiviral drug that either prevents infection or stops the production of a virus is explored. First, the efficacy of the drug is considered to decreases monotonously as time progresses. In this case, our result depicts that when the efficacy of the drug is low, the viral load decreases and increases back in time revealing the effect of the antiviral drugs is short-lived. On the other hand, for the antiviral drug with high efficacy, the viral load, as well as the number of infected cells, monotonously decreases while the number of uninfected cells increases. The dependence of the critical drug efficacy on time is also explored. Moreover, the correlation between viral load, the antiviral drug, and CTL response is also explored. In this case, not only the dependence for the basic reproduction ratio on the model parameters is explored but also we analyze the critical drug efficacy as a function of time. We show that the term related to the basic reproduction ratio increases when the CTL response step up. A simple analytically solvable mathematical model is also presented to analyze the correlation between viral load and antiviral drugs.

## I. INTRODUCTION

Viruses are tiny particles that occupy the world and have property between living and non-living things. As they are not capable of reproducing, they rely on the host cells to replicate themselves. To gain access, the virus first binds and intrudes into the host cells. Once the virus is inside the cell, it releases its genetic materials into the host cells. It then starts manipulating the cell to multiply its viral genome. Once the viral protein is produced and assembled, the new virus leaves the cell in search of other host cells. Some viruses can also stay in the host cells for a long time as a latent or chronic state. The genetic information of a virus is stored either in form of RNA or DNA. Depending on the type of virus, the host cell type also varies. For instance, the Human Immunodeficiency Virus (HIV) (see Fig. 1 [1]) directly affects Lymphocytes. Lymphocytes can be categorized into two main categories: the B and T cells. The B cells directly kill the virus by producing a specific antibody. T cells on the other hand can be categorized as killer cells (CD 8) and helper cells (CD 4). Contrary to CD 8, CD 4 gives only warning so that the cells such as CD 8 and B cells can directly kill the virus [2, 3]. Although most of these viruses contain DNA as genetic material, retroviruses such as HIV store their genetic materials as RNA. These viruses translate their RNA into DNA using an enzyme called reverse transcriptase during their life cycle. In the case of HIV, once HIV infects the patient, a higher viral load follows for the first few weeks, and then its replication becomes steady for several years. As a result, the CD 4 (which is the host cell for HIV) decreases considerably. When the CD 4 cells are below a certain threshold, the patient develops AIDS.

**FIG. 1:**
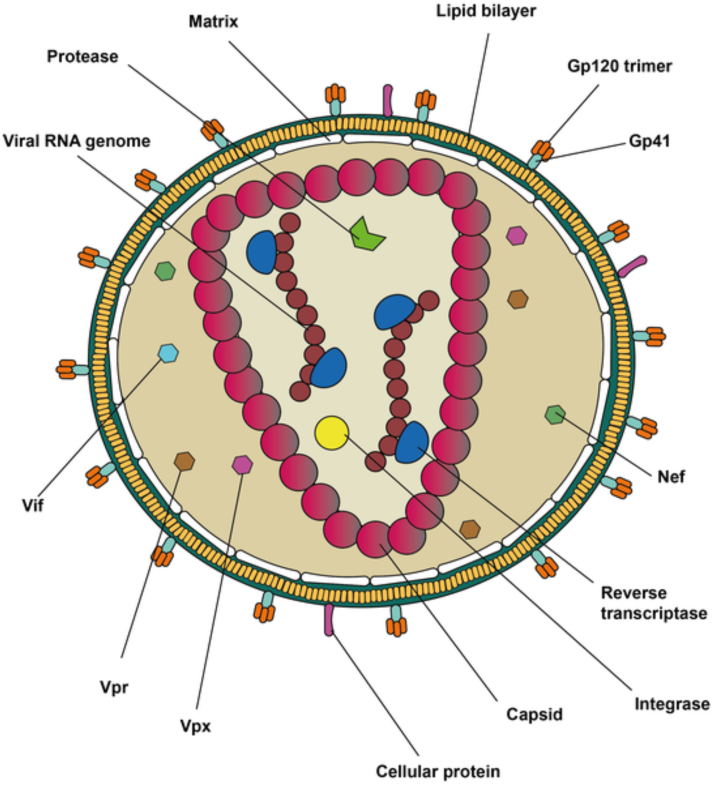
(Color online) Schematic diagram for HIV virion [1].

To tackle the spread of virulent viruses such as HIV, discovering potent antiviral drugs is vital. However, developing antiviral drugs is an exigent task since viruses mutate to overcome the effect of antiviral drugs because of this only a few antiviral drugs are currently available. Most of these drugs are developed to cure HIV and herpes virus. The fact that viruses are obligate parasites of the cells makes drug discovery complicated since the drug’s adverse effects directly affect the host cells. Many medically important viruses are also virulent and hence they cannot be propagated or tested via animal models. This in turn forces the drug discovery to be lengthy. Moreover, unlike other antimicrobial drugs, antiviral drugs have to be 100 percent potent to completely avoid drug resistance. In other words, if the drug partially inhibits the replication of the virus, through time, the number of resistant viruses will dominate the cell culture. All of the above factors significantly hinder drug discovery. Furthermore, even a potent antiviral drug does not guarantee a cure if the acute infection is already established.

Understanding the dynamics of the virus in vivo or vitro is crucial since viral diseases are the main global health concern. For instance, recent outbreaks of viral diseases such as COVID-19 not only cost trillion dollars but also killed more than 217,721 people in the USA alone. To control such a global pandemic, developing an effective therapeutic strategy is vital. Particularly, in the case of virulent viruses, mathematical modeling along with the antiviral drug helps to understand the dynamics of the virus in vivo [4]. The pioneering mathematical models on the Human Immunodeficiency virus depicted in the works [5–14] shed light regarding the host-virus correlation. Latter these model systems are modified by Neumann *et. al*. [5, 15] to study the dynamics of HCV during treatment. To study the dependence of uninfected cells, infected cells, and virus load on model parameters, Neumann proposed three differential equations. More recently, to explore the observed HCV RNA profile during treatment, Dahari *et. al*. [5, 16] extended the original Neumann model. Their findings disclose that critical drug efficacy plays a critical role. When the efficacy is greater than the critical value, the HCV will be cleared. On the contrary, when the efficacy of the drug is below the critical threshold, the virus keeps infecting the target cells.

As discussed before, the effect of antiviral drugs is short-lived since the virus mutates during the course of treatment. To include this effect, we modify Neumanny and Dahariy models. Considering the fact that the efficacy of the antiviral drug decreases in time, we rewrite the three differential equations introduced in the previous model systems. The mathematical model presented in this work analyzes the effect of an antiviral drug that either prevents infection (*e*_*k*_) or stops the production of virus (*e*_*p*_). First, we consider a case where the efficacy of the drug decreases to zero as time progresses and we then discuss the case where the efficacy of the drug decreases to a constant value as time evolves maintaining the relation 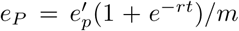 and 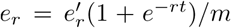. Here *r*, 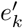 and 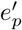 measure the ability of antiviral drug to overcome drug resistance. When *r* tends to increase, the efficacy of the drug decreases. The results obtained in this work depict that for large *r*, the viral load decreases and increases back as the antiviral drug is administered showing the effect of antiviral drugs is short-lived. On the other hand, for small *r*, the viral load, as well as the number of infected cells monotonously decreases while the host cell increases. The dependence of the critical drug efficacy on time is also explored.

The correlation between viral load, antiviral therapy, and cytotoxic lymphocyte immune response (CTL) is also explored. Not only the dependence for the basic reproduction ratio on the model parameters is explored but also we find the critical drug efficacy as a function of time. The basic reproduction ratio increases when the CTL response decline. When the viral load inclines, the CTL response step up. We also present a simple analytically solvable mathematical model to address the correlation between drug resistance and antiviral drugs.

The rest of the paper is organized as follows: in Section II, we explore the correlation between antiviral treatment and viral load. In Section III the relation between viral load, antiviral therapy, and the CTL immune response is examined. A simple analytically solvable mathematical model that addresses the correlation between drug resistance and viral load is presented in section IV. Section V deals with summary and conclusion.

## II. THE RELATION BETWEEN ANTIVIRAL DRUG AND VIRUS LOAD

In the last few decades, mathematical modeling along with antiviral drugs helps to develop a therapeutic strategy. The first model that describes the dynamics of host cells *x*, virus load *v*, and infected cells *y* as a function of time *t* was introduced in the works [8–12]. Accordingly, the dynamics of the host cell, infected cell, and virus is governed by

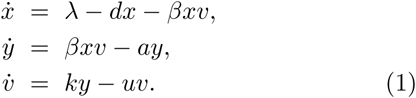

The host cells are produced at rate of *λ* and die naturally at a constant rate *d* with a half-life of 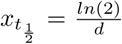. The target cells become infected at a rate of *β* and die at a rate of *a* with a corresponding half-life of 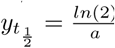. On the other hand, the viruses reproduce at a rate of *k* and die with a rate *u* with a half-life of 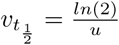 [17–20]. In this model, only the interaction between the host cells and viruses is considered neglecting other cellular activities. The host cells, the infected cells, and the viruses have a lifespan of 1*/d*, 1*/a*, and 1*/u*, respectively. During the lifespan of a cell, one infected cell produces *N* = *u/a* viruses on average [3, 19, 20].

The capacity for the virus to spread can be determined via the basic reproductive ratio

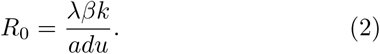

Whenever *R*_0_ *>* 1, the virus spreads while when *R*_0_ *<* 1 the virus will be cleared by the host immune system [3].

To examine the dependence of uninfected cells, infected cells and virus load on the system parameters during antiviral treatment, the above model system (Eq. (1)) was modified by Neumann *et. al*. [15] and Dahari *et. al*. [16]. The modified mathematical model presented in those works analyzes the effect of antiviral drugs that either prevents infection of new cells (*e*_*k*_) or stops production of the virion (*e*_*p*_). In this case, the above equation can be remodified to include the effect of antiviral drugs as

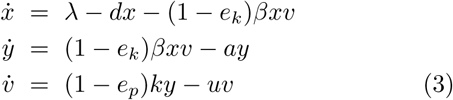

where the terms *e*_*k*_ and *e*_*p*_ are used when the antivirus blocks infection and virion production, respectively. For instance, *e*_*p*_ = 0.8 indicates that the drug has efficacy in blocking virus production by 80 percent. The antiviral drug such as protease inhibitor inhibits the infected cell from producing the right gag protein as a result the virus becomes noninfectious. A drug such as a reverse transcriptase inhibitor prohibits the infection of new cells.

Moreover, the results obtained in the last few decades depict that, usually HIV patient shows high viral load in the first few weeks of infection. As a result, the viral load becomes the highest then it starts declining for a few weeks. The viruses then keep replicating for many years until the patient develops AIDS. Since virus replication is prone to errors, the virus often develops drug resistance; HIV mutates to become drug-resistant. Particularly when the antiviral drug is administered individually, the ability of the virus to develop drug resistance steps up. However, a triple-drug therapy which includes one protease inhibitor combined with two reverse transcriptase inhibitors helps to reduce the viral load for many years [3].

Since the antiviral drugs are sensitive to time, to include this effect, next, we will modify the Neumann and Dahari model.

*Case one*.*—*As discussed before, the effect of antiviral drugs is short-lived since the virus mutates once the drug is administrated. To include this effect, we modify the above equation by assuming that 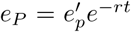 and 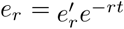. The efficacy of the drugs declines exponentially as time progresses. The decaying rate aggravates when *r* tends to increase. Hence let us rewrite Eq. (3) as

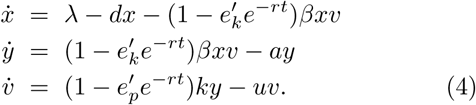

The term related to the reproductive ratio is given as

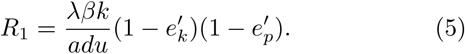

When *R*_1_ *<* 1, the antivirus drug is capable of clearing the virus and if *R*_1_ *>* 1, the virus tends to spread. At steady state,

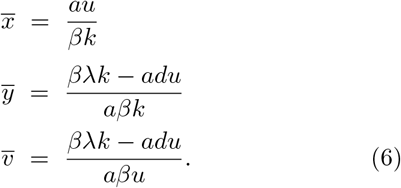

As one can note that, when *t* → ∞, *e*_*p*_ → 0 and *e*_*k*_ → 0.

At steady state, only one newly infected cell arises from one infected cell [3].

Let us next explore how the number of host cells *x*, the number of infected cells *y*, and the viral load *v* behave as a function of time by exploiting Eq. (4) numerically. From now on, all of the physiological parameters are considered to vary per unit time (days). Figure 1 depicts the plot of the number of host cells *x*, the number of infected cells *y* and the number of virus as function of time (days) for parameter choice of *λ* = 10^7^, *d* = 0.1, *a* = 0.5, *β* = 2*X*10^−9^, *k* = 1000.0, *u* = 5.0, *e*_*p*_ = 0.5, *e*_*k*_ = 0.5 and *r* = 0.06. The figure depicts that in the presence of an antiviral drug, the number of *CD*_4_ cells increases and attains a maximum value. The cell numbers then decrease and saturate to a constant value. The number of infected cells decreases and saturates to a constant value. On the other hand, the viral load decreases as the antiviral takes an effect. However, this effect is short-lived since the the viral load increases back as the viruses develop drug resistance.

When *r* is small, the ability of the antiviral drug to overcome drug resistance increases. As depicted in Fig. (2), for very small *r*, the host cells increase in time, and at a steady state, the cells saturate to a constant value. On the contrary, the infected cells as well as the plasma virus load monotonously decrease as time progresses. The figure is plotted by fixing *λ* = 10^7^, *d* = 0.1, *a* = 0.5, *β* = 2*X*10^−9^, *k* = 1000, *u* = 5.0, *e*_*p*_ = 0.9, *e*_*k*_ = 0.9 and *r* = 0.0001. These all results indicate that when combined drugs are administrated, the viral load is significantly reduced depending on the initial viral load. Figure 3 is plotted by fixing *λ* = 10^7^, *d* = 0.1, *a* = 0.5, *β* = 2*X*10^−9^, *k* = 1000, *u* = 5.0, *e*_*p*_ = 0.9, *e*_*k*_ = 0.9 and *r* = 20.0. The figure exhibits that for large *r*, the *CD*_4_ cells decrease and exhibit a local minima. As time progresses, the number of cells increases and saturates to a constant value. On the other hand, the number of infected cells *y* and the viral load *v* decreases and saturates to considerably large value as time progresses. Figure 3 also depicts that when the drug is unable to control the infection in a very short period of time, the number of drug resistant viruses steps up.

**FIG. 2:**
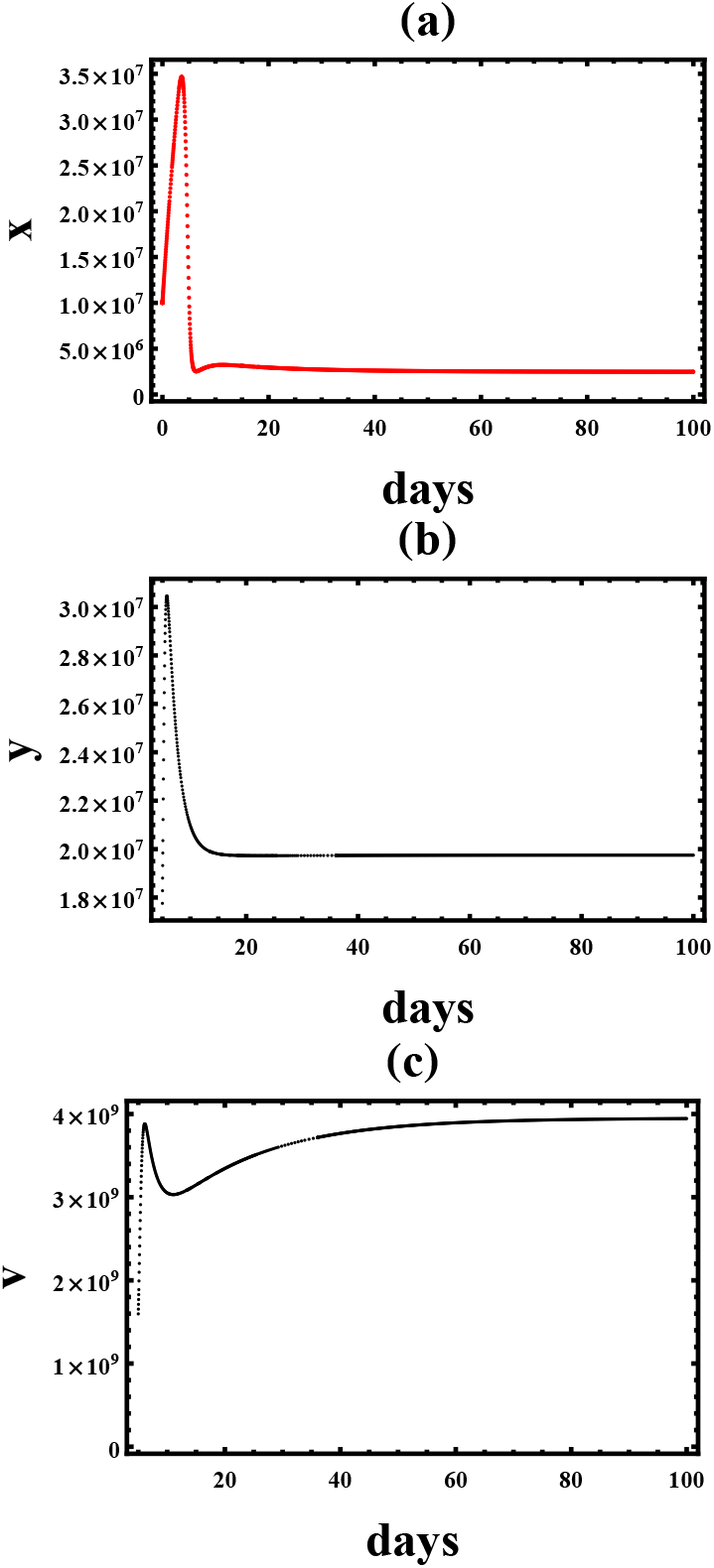
(Color online) (a) The number of host cells *x* as a function of time (days). (b) The number of infected cells as function of time (days). (c) The virus load as a function of the time (days). In the figure, we fix *λ* = 10^7^, *d* = 0.1, *a* = 0.5, *β* = 2*X*10^−9^, *k* = 1000.0, *u* = 5.0, *e*_*p*_ = 0.5, *e*_*k*_ = 0.5 and *r* = 0.06.

**FIG. 3:**
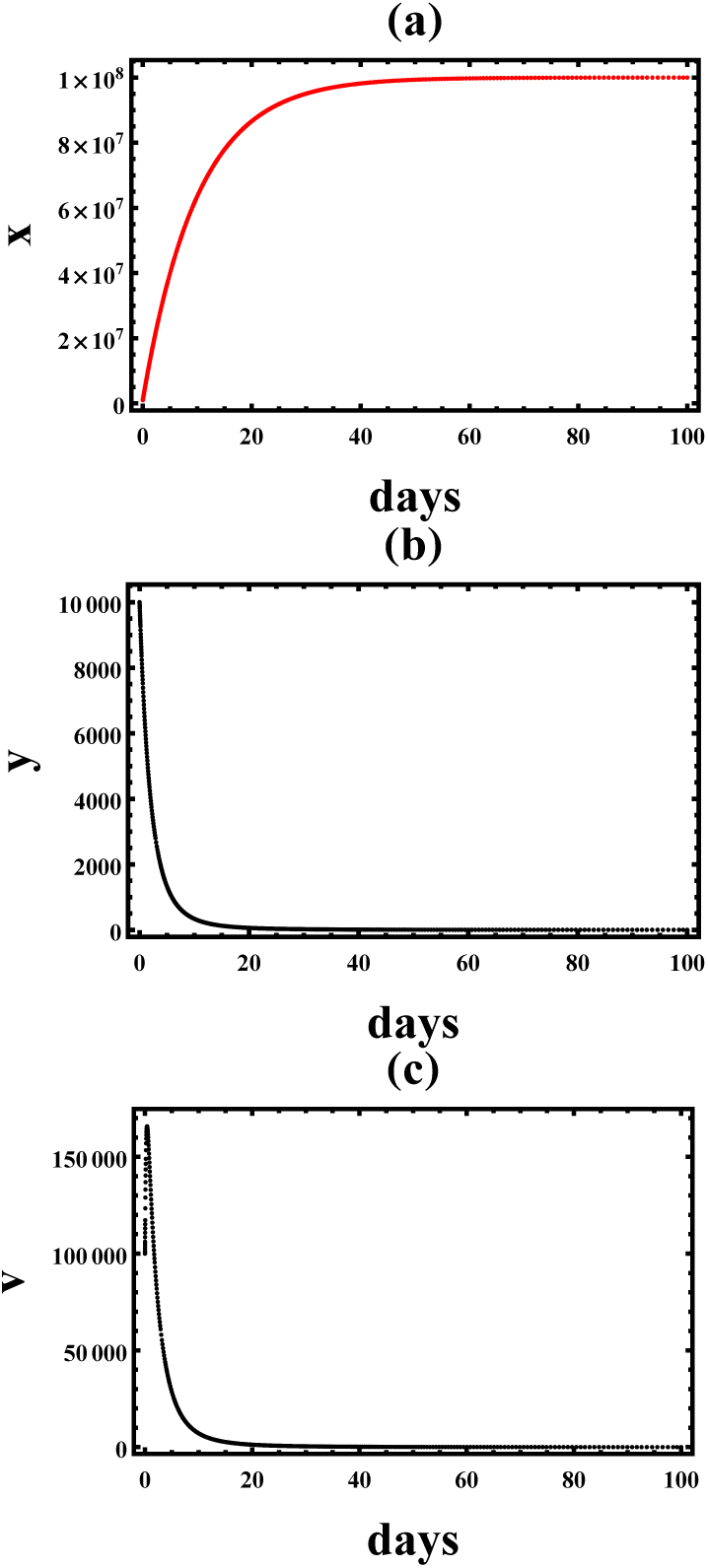
(Color online) (a) The number of host cells *x* as a function of time (days). (b) The number of infected cells as function of time (days). (c) The plasma virus load as a function of time (days). In the figure, we fix *λ* = 10^7^, *d* = 0.1, *a* = 0.5, *β* = 2*X*10^−9^, *k* = 1000.0, *u* = 5.0, *e*_*p*_ = 0.9, *e*_*k*_ = 0.9 and *r* = 0.0001.

**FIG. 4:**
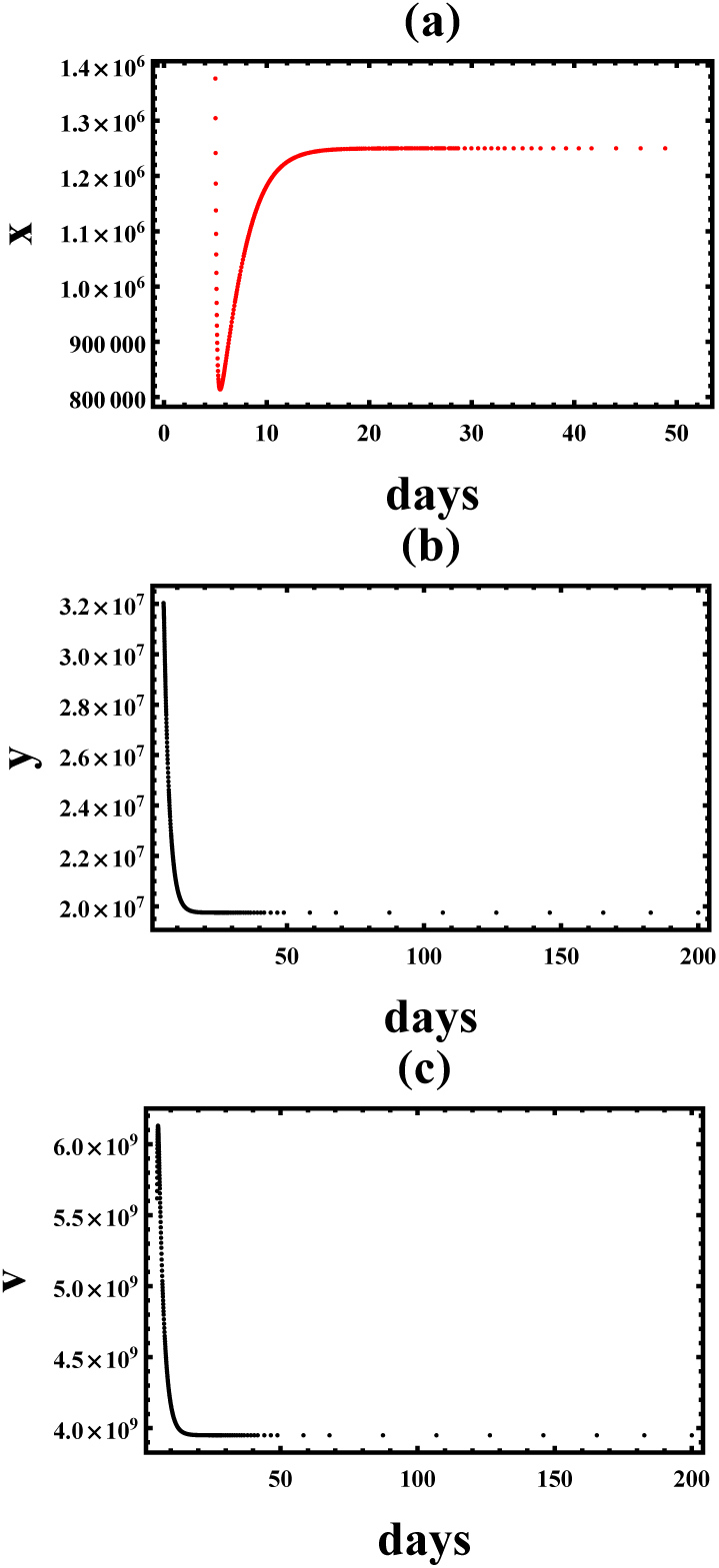
(Color online) (a) The number of host cells *x* as a function of days. (b) The number of infected cells as function of days. (c) The virus load as a function of the days. In the figure, we fix *λ* = 10^7^, *d* = 0.1, *a* = 0.5, *β* = 2*X*10^−9^, *k* = 1000.0, *u* = 5.0, *e*_*p*_ = 0.9, *e*_*k*_ = 0.9 and *r* = 20.0.

*Case two*.*—* In the previous case, the efficacy of the drug is considered to decrease monotonously as time progresses. In this section, the efficacy of the drug is assumed to decrease to a constant value as time increases maintaining the relation 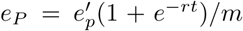 and 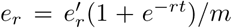. The dynamics of host cells, infected cells, and viral load is governed by the equation

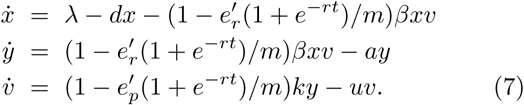

After some algebra, the term related to the basic reproductive ratio reduces to

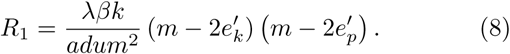

As one can see from Eq. (8) that as *m* steps up, the drug losses its potency and as a result *R*_1_ increases. When *R*_1_ *<* 1, the antivirus drug treatment is successful and this occurs for large values of *e*_*p*_ and *e*_*k*_. When *R*_1_ *>* 1, the virus overcomes the antivirus treatment.

At equilibrium, one finds

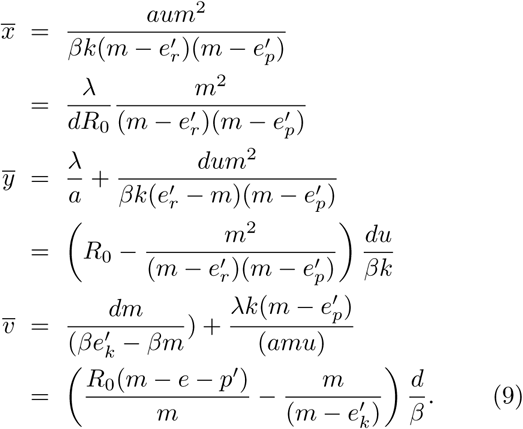

The case *R*_0_ ≫ 1 indicates that the equilibrium abundance of the uninfected cells is much less than the number of uninfected cells before treatment. When the drug is successful, (large values of 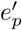 or 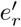), the equilibrium abundance of the uninfected cells increases. On the contrary, for a highly cytopathic virus (*R*_1_ ≫ 1), the number of infected cells, as well as the viral load steps up. When 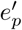 and 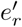 increase, the equilibrium abundance of infected cells as well as viral load decreases.

In general for large *R*_0_, Eq. (9) converges to

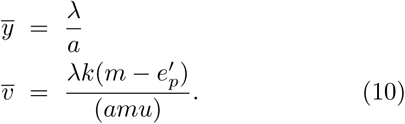

Clearly 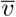 decreases as 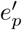 and 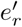 increase.

The overall efficacy can be written as [16]

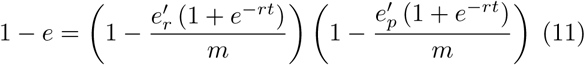

where 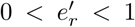 and 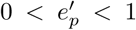. At steady state 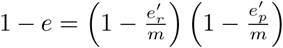. The transcritical bifurcation point (at steady state) can be analyzed via Eq. (9) and after some algebra we find

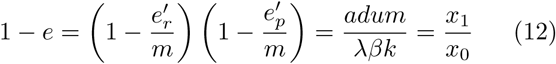

where 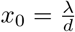 denotes the number of uninfected host cells before infection and 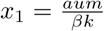 designates the number of uninfected cells in the chronic case. This implies the critical efficacy is given as 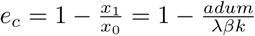.

To write the overall efficacy as a function of time, for simplicity, let us further assume that 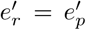. In this case, Eq. (12) can be rewritten as

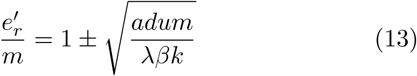

and hence

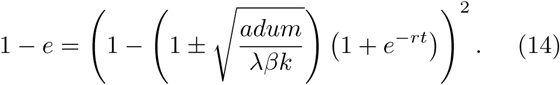

From Eq. (14), one finds

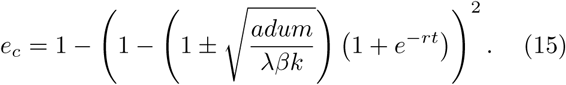

The critical efficacy serves as an alternative way of determining whether antiviral treatment is successful or not. When *e > e*_*c*_, the antiviral clears the infection and if *e < e*_*c*_, the virus replicates.

## III. THE CORRELATION BETWEEN ANTIVIRAL DRUG, IMMUNE RESPONSE AND VIRUS LOAD

The basic mathematical model that specifies the relation between the immune response, antiviral drug, and viral load is given by

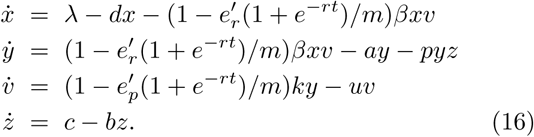

Once again the terms *x, y*, and *v* denote the uninfected cells, infected cells, and the viral load. The term *z* denotes the CTL response and the CTL die at a rate of *b* and produced at a rate of *c*. The term CTL is defined as cytotoxic lymphocytes that have responsibility for killing the infected cells. The term related to the basic reproduction rate is given as

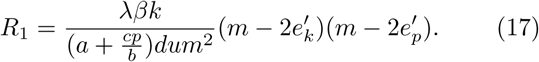

It vital to see that when *R*_0_ *>* 0 the virus becomes successful to persist an infection which triggers an immune responses *z*. As long as the coordination between the immune response and the antiviral drug treatment is strong enough, the virus will be cleared *R*_1_ *<* 1. As it can be clearly seen from Eq. (17), when the CTL response step up, *R*_1_ declines as expected.

The equilibrium abundance of the host cells, infected cells, viral load, and CTL response can be given as

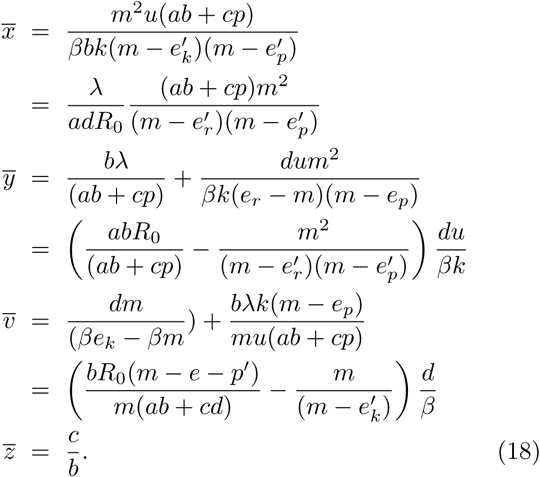

Exploiting Eq. (18), one can deduce when *R*_0_ ≫ 1, the equilibrium abundance of the uninfected cells becomes much lower in comparison to the number of cells before treatment. The equilibrium abundance of the uninfected cells steps up when there is CTL response (when *c* and *p* increase) or when the antiviral treatment is successful (when 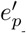 and 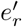 increase). On the contrary, 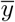 and 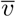 decline whenever there is a strong CTL response or when the antiviral treatment is successful.

In Fig. 5a, we plot the phase diagram for a regime *R*_1_ *<* 1 (shaded region) in the phase space of *e*_*k*_ and *e*_*p*_. In Figure 5b, the phase diagram for a regime *R*_1_ *<* 1 (shaded region) in the phase space of *m* and *e*_*p*_ = *e*_*k*_ is plotted. In the figure, we fix *λ* = 10^7^, *d* = 0.1, *a* = 0.5, *β* = 2*X*10^−9^, *k* = 1000, *u* = 5, *m* = 2, *r* = 0.0001, *p* = 1.0, *b* = 0.5 and *c* = 2.0.

**FIG. 5:**
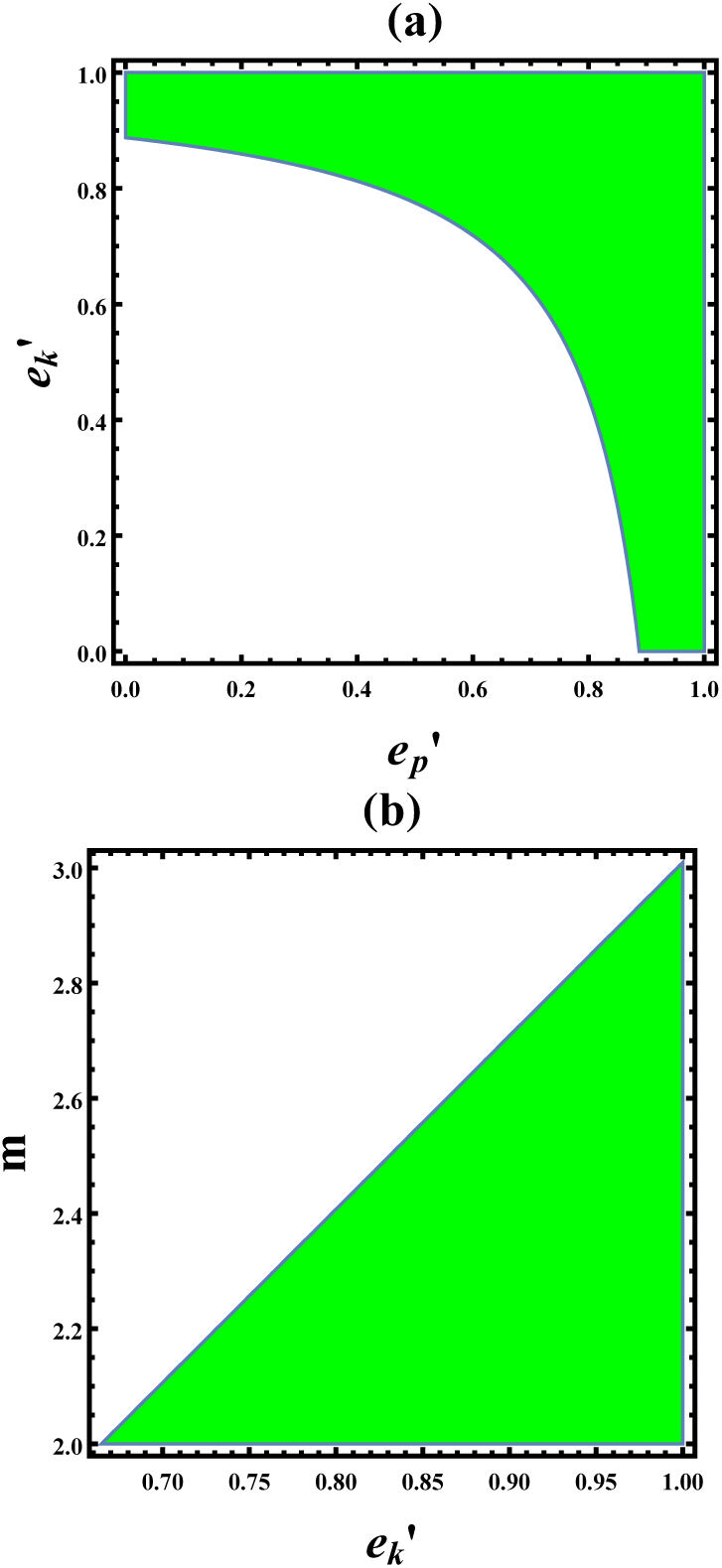
(Color online) (a) The phase diagram for a regime *R*_0_ *<* 1 in the phase space of *e*_*k*_ and *e*_*p*_. (b) The phase diagram for a regime *R*_0_ *<* 1 in the phase space of *m* and *e*_*p*_ = *e*_*k*_. In the figure, we fix *λ* = 10^7^, *d* = 0.1, *a* = 0.5, *β* = 2*X*10^−9^, *k* = 1000, *u* = 5, *m* = 2, *r* = 0.0001, *p* = 1.0, *b* = 0.5 and *c* = 2.0.

Furthermore, the transcritical bifurcation point can be analyzed via Eq. (18) and after some algebra (at steady state) we find

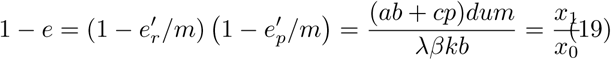

where 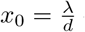 denotes the number of uninfected host cells before infection and 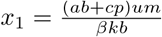 designates the number of uninfected cells in a chronic case. This implies the critical efficacy is given as 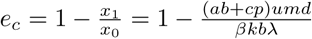.

Assuming 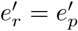, Eq. (19) can be rewritten as

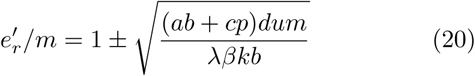

and the overall efficacy as a function of time is given by

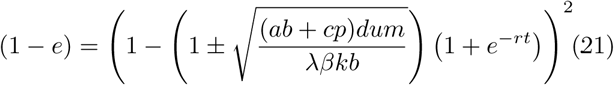

From Eq. (21), one finds

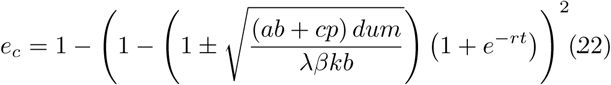

Once again, the infection will be cleared when *e > e*_*c*_, and the virus replicates as long as *e < e*_*c*_.

In order to get a clear insight, let us explore Eq. (16). In Fig. 6, the number of host cells *x*, the number of infected cells and the viral load as a function of time is plotted. In the figure, we fix *λ* = 10^7^, *d* = 0.1, *a* = 0.5, *β* = 2*X*10^−9^, *k* = 1000, 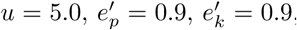, *r* = 0.1, *p* = 10.0, *b* = 0.5 and *c* = 1.0. For such parameter choice, *R*_0_ ≫ 1.0 and *R*_1_ ≪ 1.0 revealing that initially, the virus establishes an infection but latter the antiviral drug and CTL response collaborate to clear the infection. As a result, the number of host cells increases while the viral load as well as the infected cells decreases. Since the virus is responsible for initiating the CTL response, as the viral load declines, the CTL response step down.

**FIG. 6:**
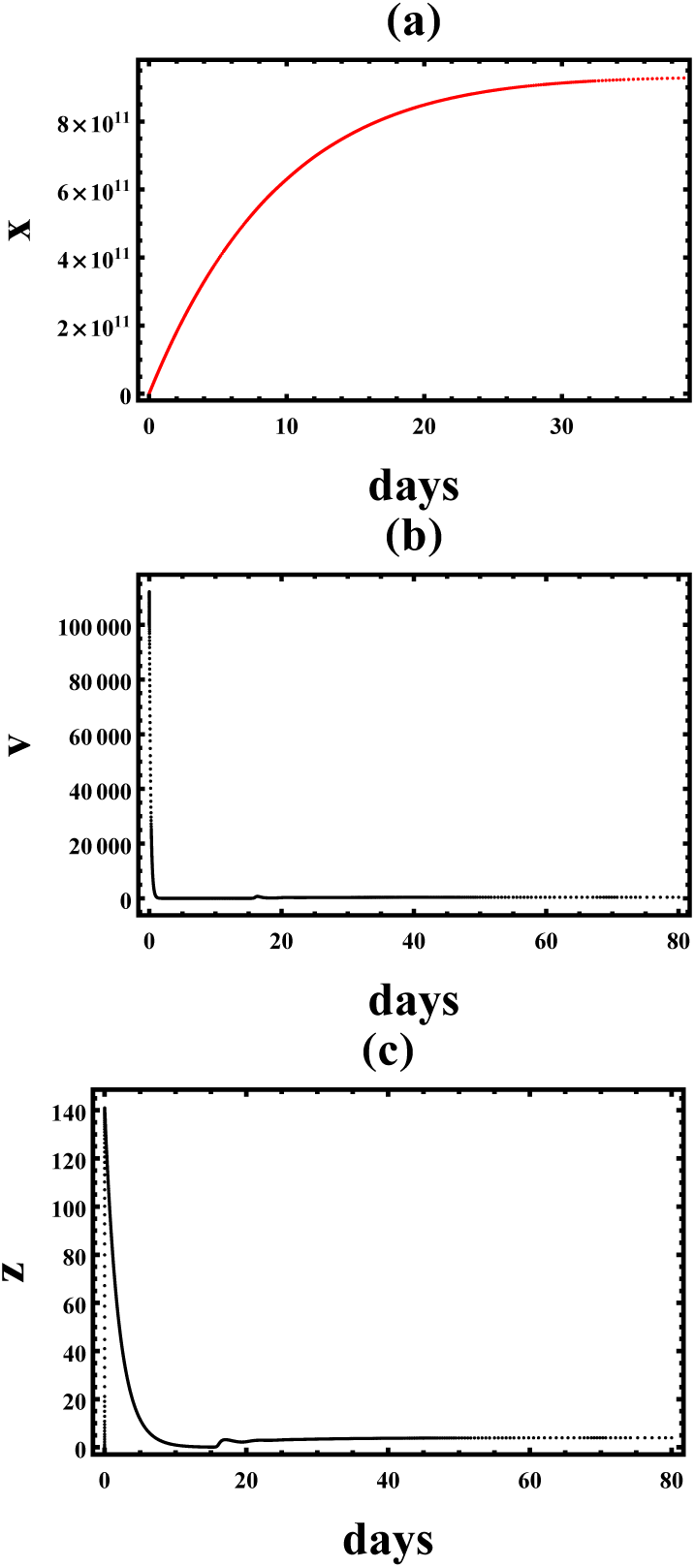
(Color online) (a) The number of host cells *x* as a function of days. (b) The number of infected cells as function of days (c) The virus load as a function of the days. In the figure, we fix *λ* = 10^7^, *d* = 0.1, *a* = 0.5, *β* = 2*X*10^−9^, *k* = 1000, *u* = 5.0, 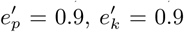, *r* = 0.1, *p* = 10.0, *b* = 0.5 and *c* = 1.0.

**FIG. 7:**
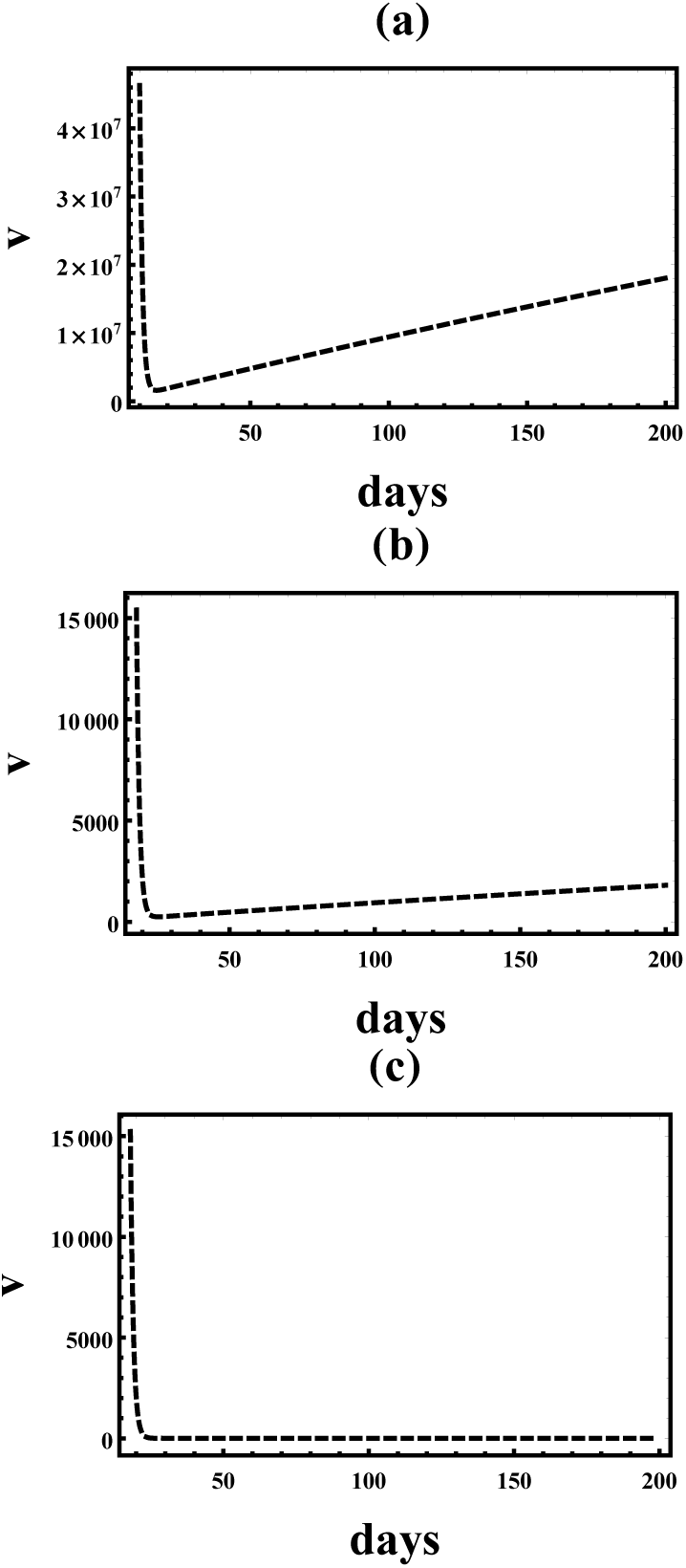
(Color online) The virus load as a function of time. In the figure, we fix *u* = 1.0. (a) Single drug therapy *s* = 1. (b) Double drug therapy *s* = 2. (c) Triple drug therapy *s* = 3.

## IV. THE DYNAMICS OF MUTANT VIRUS IN THE PRESENCE OF ANTIVIRAL DRUGS

As discussed before, the fact that viruses are an obligate parasite of the cells forces drug discovery to be complicated as the drug’s adverse effects directly affect the host cells. Many medically important viruses are also virulent and hence they cannot be propagated or tested via animal models. This in turn forces the drug discovery to be lengthy. Moreover, unlike other antimicrobial drugs, antiviral drugs have to be 100 percent potent to completely avoid drug resistance. In other words, if the drug partially inhibits the replication of the virus, through time, the number of the resistant virus will dominate the cell culture.

To discuss the dynamics of the mutant virus, let us assume that the virus mutates (when a single drug is administered) by changing one base every 10^4^ viruses. If 10^11^ viruses are produced per day, then this results in 10^7^ mutant viruses. On the contrary, when two antiviral drugs are administrated, 10^3^ mutant viruses will be produced. In the case of triple drug therapy, no mutant virus is produced. To account for this effect, let us remodify the rate of mutant virus production per day as

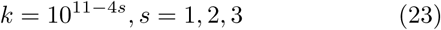

where the variable *s* = 1, 2, 3 corresponds to a single, double, and triple drug therapy, respectively.

Here we assume only the rate of mutant virus production *k* determines the dynamics and 10^11^ viruses are produced per day. To get an instructive analytical solution regarding the relation between antiviral drug and viral load, let us solve the differential equation

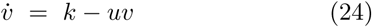

neglecting the effect of uninfected and infected host cells. Here the mutant virus produced at rate of *k* and die with the rate of *u*. The solution for the above equation is given as

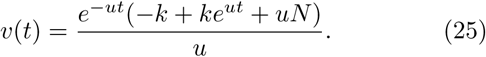

Whenever 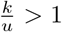 the virus spreads and when 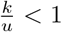, the antiviral is capable of eliminating the virus.

Exploiting Eq. (25) one can comprehend that, in the case of single therapy, the virus load decreases during the course of treatment. As time progresses, the viral load increases back due to the emergence of drug resistance (see Fig. 8a). In the case of double drug therapy, as shown in Fig. 8b, the viral load decreases but relapses back as time progresses. When triple drugs are administered, the viral replication becomes suppressed as depicted in Fig 8c. The readers should understand that the triple drug therapy does not guarantee a cure. If the patient halts his or her therapy, the viral replications will resume because of the latent and chronic infected cells. At steady state, we get

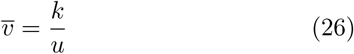

At equilibrium, the viral load spikes as *k* increases and it decreases as *u* steps up.

## V. SUMMARY AND CONCLUSION

Developing antiviral drugs is challenging but an urgent task since outbreaks of viral diseases not only killed several people but also cost trillion dollars worldwide. The discovery of new antiviral drugs together with emerging mathematical models helps to understand the dynamics of the virus in vivo. For instance, the pioneering mathematical models on HIV shown in the works [8–14] disclose the host-virus correlation. Moreover, to study the correlation between, antiviral drugs and viral load, an elegant mathematical model was presented in the works [15, 16].

Due to the emergence of drug resistance, the efficiency of antiviral drugs is short-lived. To study this effect, we numerically study the dynamics of the host cells and viral load in the presence of an antiviral drug that either prevents infection (*e*_*k*_) or stops the production of virus (*e*_*p*_). For the drug whose efficacy depends on time, we show that when the efficacy of the drug is low, the viral load decreases and increases back in time revealing the effect of the antiviral drugs is short-lived. On the contrary, for the antiviral drug with high efficacy, the viral load, as well as the number of infected cells, monotonously decreases while the number of uninfected cells increases. The dynamics of critical drug efficacy on time is also explored. Furthermore, the correlation between viral load, an antiviral drug, and CTL response is also explored. Not only the dependence for the basic reproduction ratio on the model parameters is explored but also we analyze the critical drug efficacy as a function of time. The term related to the basic reproduction ratio increases when the CTL response step up. A simple analytically solvable mathematical model to analyze the correlation between viral load and antiviral drugs is also presented.

In conclusion, in this work, we present a simple model which not only serves as a basic tool for better understanding of viral dynamics in vivo and vitro but also helps in developing an effective therapeutic strategy.

## Acknowledgment

I would like to thank Mulu Zebene and Blaynesh Bezabih for the constant encouragement.

## References

[1] This file is made available by MechESR under the Creative Commons CC 1.0 Universal Public Domain Dedication.Images (Wikimedia Commons).

[2] G. Doitsh and W. C. Greene, Cell Host and Microbe 19, 280 (2016).

[3] Nowak, M. A., and May, R. (2001). Virus Dynamics: Mathematical Principles of Immunology and Virology. Oxford University Press.

[4] S. Wang, Y. Pan, Q. Wang, H. Miao, A. N. Brown and L. Rong, Mathematical Biosciences 328, 108438 (2020).

[5] C. Zitzmann and L. Kaderali, Forntiers in Microbiology 9, 1546 (2018).

[6] Y. Wang, J. Liu and L. Liu, Advances in Difference Equations 1, 225 (2018).

[7] S. S. Chen, C. Y. Cheng and Y. Takeuchi, Journal of Mathematical Analysis and Applications 442, 642 (2016).

[8] A. S. Perelson, D. E. Kirschner and R. De Boer, Math. Biosci 114, 81 (1993).

[9] A. S. Perelson,, A. U. Neumann, M. Markowitz, J. M Leonard and D. D Ho, Science 271, 1582 (1996).

[10] A. S. Perelson, P. Essunger, Y. Cao, M. Vesanen, A. Hurley and K. Saksela, Nature 387, 188 (1997).

[11] Ho, D. D., Neumann, A. U., Perelson, A. S., Chen, W., Leonard, J. M., and Markowitz, M. (1995). Rapid turnover of plasma virions and CD4 lymphocytes in HIV-1 infection. Nature 373, 123126. doi: 10.1038/373123a.

[12] Bonhoeffer, S., May, R. M., Shaw, G. M., and Nowak, M. A. (1997). Virus dynamics and drug therapy. Proc. Natl. Acad. Sci. U.S.A. 94, 69716976. doi: 10.1073/pnas.94.13.6971.

[13] Stafford, M. A., Corey, L., Cao, Y., Daar, E. S., Ho, D. D., and Perelson, A. S. (2000). Modeling plasma virus concentration during primary HIV infection. J. Theor. Biol. 203, 285301. doi: 10.1006/jtbi.2000.1076

[14] Wei, X., Ghosh, S. K., Taylor, M. E., Johnson, V. A., Emini, E. A., Deutsch, P., et al. (1995). Viral dynamics in human immunodeficiency virus type 1 infection. Nature 373, 117122. doi: 10.1038/373117a0

[15] Neumann, A. U. (1998). Hepatitis C viral dynamics in vivo and the antiviral efficacy of interferon-therapy. Science 282, 103107. doi: 10.1126/science.282.5386.103

[16] Dahari, H., Lo, A., Ribeiro, R. M., and Perelson, A. S. (2007a). Modeling hepatitis C virus dynamics: Liver regeneration and critical drug efficacy. J. Theor. Biol. 247, 371381. doi: 10.1016/j.jtbi.2007.03.006

[17] Nowak, M. A., and Bangham, C. R. (1996). Population dynamics of immune responses to persistent viruses. Science 272, 7479.

[18] Nowak, M. A., Bonhoeffer, S., Hill, A.M., Boehme, R., Thomas, H. C., andMcDade, H. (1996). Viral dynamics in hepatitis B virus infection. Proc. Natl. Acad. Sci. U.S.A. 93, 43984402. doi: 10.1073/pnas.93.9.4398

[19] Perelson, A. S. (2002). Modelling viral and immune system dynamics. Nat. Rev. Immunol. 2, 2836. doi: 10.1038/nri700

[20] Wodarz, D., and Nowak, M. A. (2002).Mathematical models of HIV pathogenesis and treatment. BioEssays 24, 11781187. doi: 10.1002/bies.10196

